# Patterning of a telencephalon-like region in the adult brain of amphioxus

**DOI:** 10.1101/307629

**Authors:** Èlia Benito-Gutiérrez, Manuel Stemmer, Silvia D Rohr, Laura N Schuhmacher, Jocelyn Tang, Aleksandra Marconi, Gáspár Jékely, Detlev Arendt

## Abstract

The evolutionary origin of the vertebrate telencephalon remains unsolved. A major challenge has been the identification of homologous brain parts in invertebrate chordates. Here we report evidence for a telencephalic region in the brain of amphioxus, the most basally branching invertebrate chordate. This region is characterised, like its vertebrate counterpart, by the combined expression of the telencephalic markers *FoxG1, Emx* and *Lhx2/9*. It is located at the anterior neural border and dorsal-ventrally patterned, as in vertebrates, by the antagonistic expression of *Pax4/6* and *Nkx2.1*, and a ventral *Hh* signal. This part of the brain develops only after metamorphosis via sustained proliferation of neuronal progenitors at the ventricular zone. This is concomitant with a massive expansion of late differentiating neuronal types as revealed by neuropeptide and neurotransmitter profiling. Overall, our results suggest that the adult amphioxus brain shows remarkable similarities to the vertebrate embryonic brain, thus providing a key missing link in understanding the invertebrate-to-vertebrate transition in chordate brain evolution.

## INTRODUCTION

Vertebrates, unlike most other animals, have a dorsal tubular brain. This develops from the anterior portion of the neural tube, which forms three primary vesicles (forebrain (telencephalon + diencephalon), midbrain and hindbrain). These vesicles progressively sub-regionalise, forming the different brain regions around an also growing ventricular system. Molecularly, this involves the patterning of regions by specific sets of genes that are induced by organising centres (Tole and Hebert 2013). No evidence of vertebrate brain-like divisions or brain-like organisers have yet been found in the invertebrate chordates, amphioxus and ascidians, which already possess a tubular nervous system. For example, the specific telencephalic marker *FoxG1* (*Bf-1*) is only sporadically expressed in the amphioxus brain, while *Emx* expression has not been analysed yet (Toresson et al., 1998; Williams and Holland 2000). Similarly, in ascidians, *Emx* is not expressed in the central nervous system (Oda and Saiga, 2001) and *FoxG1* expression has not yet been described. Consequently, in the context of vertebrate brain evolution the telencephalon has been long regarded as a vertebrate innovation (Northcutt 2005). More recently, however, the identification of specific cells with telencephalic or boundary-organiser profiles in non-chordate invertebrates has indicated that, alternatively, invertebrate chordates may have lost parts of their initial brain complexity (Tomer et al., 2010; Pani et al., 2013).

We herein demonstrate the presence of a telencephalon-like region in the brain of amphioxus. Crucially, we find this region only in adult animals, the molecular profile of which has never before been studied. Previous ultrastructural studies of the adult amphioxus brain had revealed that the organisation of the brain dramatically changes after metamorphosis, displaying a different anatomy with respect to that of the embryo (Wicht and Lacalli 2005). In accordance with these descriptions we observe that a large portion of the dorsal and anterior brain of amphioxus develops late, revealing new domains of gene expression where telencephalic markers are enriched. We show this in whole-brain image reconstructions of serially sectioned adult brains. We focus on genes whose function is essential for telencephalon development in vertebrates, such as *FoxG1, Emx*, *Fezf* (in Suppl) and *Lhx2* (Beccari et al., 2013); and on genes whose mutation results in the specific loss of particular cell types within the vertebrate telencephalon, such as *Hh, Nkx2.1* and *Pax4/6*. We also investigate patterns of neuronal differentiation and growth and conclude that a substantial portion of brain development occurs very late, during periods of metamorphosis and settlement This late differentiation explains why previous studies on embryonic and larval stages suggested a considerably simpler brain organisation in amphioxus. Our results suggest that the ancestral chordate brain already had a regionalised ventricular system with a telencephalon-like region.

## RESULTS

### Telencephalic transcription factors are expressed in the adult amphioxus brain

We have found that in the adult amphioxus brain *FoxG1* is expressed in a broad anterior domain that occupies almost the entire cerebral vesicle (**Fig 1B-E**). This domain specifically extends from the floor of the cerebral vesicle (**Fig1B**) to a point about 14-16 microns above, before the ventricle closes dorsally by the Joseph Cells Mantle that caps the brain. This contrasts with the limited expression of *FoxG1* earlier in development, when it is only expressed by a few cells located right under the pigment of the frontal eye (**Fig 1R**). In the adult brain FoxG1 + cells show an anterior limit adjacent to the pigment of the frontal eye and a caudal limit that coincides with the anterior part of the infundibular organ (IO) (**Fig 1S**). The frontal eye and the IO have previously been compared to the vertebrate eyes and the flexural and subcomissural organs, respectively (Vopalenski et al., 2012; Gobron et al., 1999). These demarcate the anterior-posterior limits of the entire vertebrate telencephalon, thus suggesting that the expression of FoxG1 in adult amphioxus brains parallels the extensive telencephalic expression of *FoxG1* in vertebrates (Martynoga et al., 2005; Sugahara et al., 2016) (**Fig1T**) (see also *FoxG1* expression in the Allen Brain Atlas at P28). Moreover, similar to amphioxus, in mouse and zebrafish, this entire region develops from a single primary anterior vesicle, which expresses *FoxG1* at a very early stage (Martynoga et al., 2005; Danesin et al., 2009; Roth et al., 2010).

**Fig 1.**
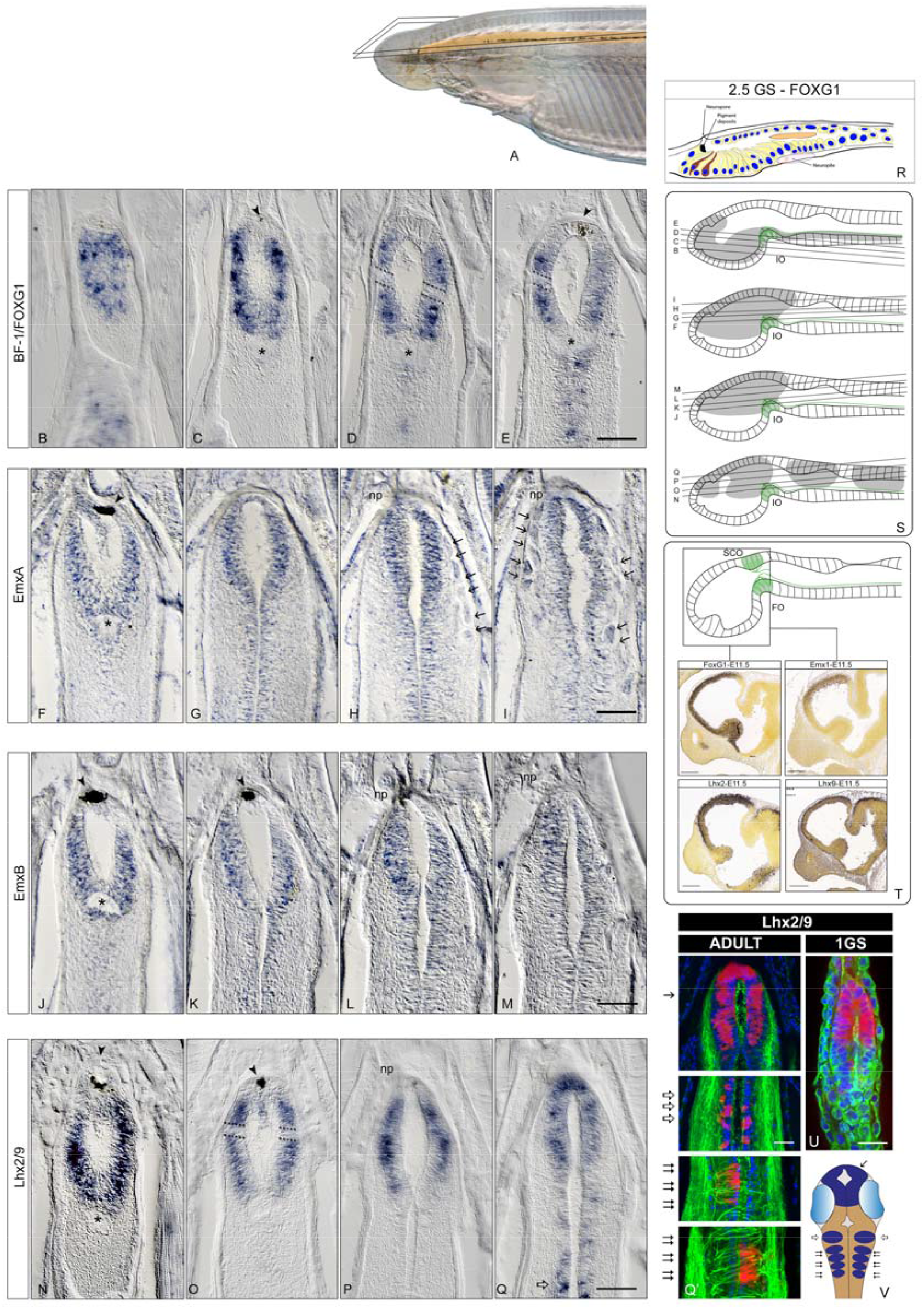
Expression of telencephalic markers in the adult brain of amphioxus. **A**. Shows the head of an amphioxus with the brain and neural tube highlighted in orange and the sectioning planes superimposed. Expression of *Bf1/FoxG1* (**B-E**), *EmxA* (**F-I**), *EmxB* (**J-M**) and *Lhx2/9* (**N-Q**) is shown in coronal paraffin sections from ventral to dorsal with the ventricle of the cerebral vesicle (cv) centred in the images. *FoxG1* expression extends throughout the ventral and lateral walls of the cv in adults (**B-E**). *EmxA* (**F-I**) and *EmxB* (**J-M**) are expressed in the adult in the dorsal three quarters of the cerebral vesicle. Only *EmxA* reaches the top mantle of Joseph cells, indicated with arrows in **H** and **I**. *Lhx2/9* expression (**N-Q**) spans throughout the entire dorsal half of the cv with a dorsal limit just below the mantle of Joseph cells. **R**. The expansion of the *FoxG1* domain in adults is particularly obvious when compared to that of larvae, where only few cells below the pigment of the frontal eye are found (brown coloured cells in a schematic representation modified after Vopalenski et al., 2012). **S**. Brain sectioning planes for *Bf1/FoxG1* (**B-E**), *EmxA* (**F-I**), *EmxB* (**J-M**) and *Lhx2/9* (**N-Q**) on a lateral representation of an adult amphioxus brain (note that the anterior-most pigment of the frontal eye and the dorsal-most Joseph Cells Mantle are not represented in the diagrams for clarity). Grey shading indicates the expression domains for each gene. **T**. Representation of a salmon embryo brain at the stage when the flexural organ (FO) and the sub-commissural organ (SCO) co-exist at the level of the cephalic flexure (modified after Nieuwenhuys et al., 2013) for comparison with the also highlighted in green infundibular organ (IO) of amphioxus in **S**. The frame in **T** indicates similar sectioning planes of the expression of *FoxG1, Emx1, Lhx2* and *Lhx9* in mice embryos at the earliest embryonic stage available in the Allen Brain Atlas (images extracted from the Allen Brain Atlas) to compare with the respective in amphioxus. **U**. Early in development *Lhx2/9* is only expressed in the anterior half of the brain of amphioxus, as shown in a confocal image of a whole mount *in situ*. This contrasts with the more complex *Lxh2/9* pattern seen in adult amphioxus, shown by confocal microscopy in **Q’** from the same section in **Q**, which recapitulates the expression pattern of both *Lhx2* and *Lhx9* early in the development of zebrafish, schematically represented in **V**: *Lxh2* shown in light blue (eyes) and overlapping *Lhx2* and *Lhx9* in dark blue (drawn based on Ando et al., 2005). Empty arrows indicate the expression of *Lhx2* and *Lhx9* at the mid-hindbrain boundary region and double arrows that at of the reticulo-spinal neurons. Confocal images in **Q’** and **U** are counterstained with acetylated tubulin (shown in green) to label the axonal scaffold and with DAPI (shown in blue) to identify the nuclei of individual cells. All scale bars in amphioxus sections are 50μm, except in **Q’** that is 20μm as in **U**. Scale bars in mice sections are 346μm. Abbreviations: 1GS: One gill slit; 2.5: Two and a half gill slits; np: neuropore. Arrowheads indicate the pigment of the frontal eye. Discontinuous lines show lateral gaps of expression at half way in the anterior-posterior axis of the cv in the FoxG1 and Lhx2/9 domains.

In mouse, the dorsal portion of the *FoxG1* domain partially overlaps with the expression of *Emx* (**Fig1T**). To explore if this was also the case in amphioxus, we examined the expression of two of the three *Emx* genes in amphioxus: *EmxA* and *EmxB* (Williams and Holland, 2000; Takatori et al., 2008). To date, no expression of *Emx* has been reported in the nervous system of amphioxus. We found that both *EmxA* and *EmxB* were expressed in the dorsal half of the adult cerebral vesicle, overlapping the dorsal part of the *FoxG1* domain, and with a rostral limit behind the frontal eye and a caudal limit coincident with the IO (**Fig1F-M**). *EmxB* shows a more restricted expression mostly overlapping with the ventral side of *EmxA*, while *EmxA* expression can also be detected in the Joseph Cells Mantle that caps the brain (**Fig1H-I and arrows**).

In vertebrates, *Lhx2* and *Lhx9* are also essential for telencephalon development. Consistent with the *FoxG1* and *EmxA/B* expression, the amphioxus Lhx2/9 pro-orthologous gene is also widely expressed in the dorsal cerebral vesicle of the adult amphioxus (**Fig1N-Q**). As in the vertebrate telencephalon, the anterior *Lhx2/9* domain shows overlapping expression with *FoxG1* and *Emx* in the dorsal half of the cerebral vesicle (**Fig1C, F, J and N**). Adding to this, the patched pattern of *Lhx2/9* in the posterior amphioxus brain matches similar domains in the zebrafish embryo, where *Lhx2* and *Lhx9* also mark the midbrain/hindbrain boundary and the bilaterally arranged reticulo-spinal neurons (Ando et al., 2005) (**Fig1V**). In amphioxus the most posterior bilaterally arranged Lhx2/9+ neurons are asymmetrically paired (**Fig 1Q-Q’**), probably following the left-right offset of the somites and associated nerve roots. For comparison, we also determined the embryonic expression of *Lhx2/9* and found it confined to the most anterior part of the cerebral vesicle only, with a caudal limit roughly at the middle of the ventricle in one-gill-slit embryos (**Fig1U**). Consequently, the tripartite expression of *Lhx2/9*, as observed in adults, is indicative of new brain compartments and boundaries forming later in development.

### The adult amphioxus brain has a dorsoventrally regionalized telencephalic domain

We next explored the expression patterns of genes with known roles in dorsoventral regionalisation within the vertebrate telencephalon. In zebrafish and mouse, the telencephalon is divided into a dorsal (pallial) and a ventral (subpallial) domain characterised by the expression of *Pax6* and *Nkx2.1*, respectively (Hebert and Fishell, 2008). The ventral limit of *Pax6* defines the subpallial-pallial boundary (Tole and Hebert 2013, Cocas et al., 2011, Sugahara et al, 2011) helped by the antagonistic interaction with Nk2.1, essential to establish the correct position of this boundary (Nat et al., 2013). *Shh* is expressed in a ventromedial domain partially overlapping *Nkx2.1* (Hebert and Fishell, 2008; Sugahara et al., 2016). It induces formation of the telencephalon by activating *FoxG1* early in vertebrate development (Retaux and Kano 2010; Fuccillo et al., 2014).

In amphioxus, *Pax4/6*, the orthologue of *Pax6* in vertebrates, has been studied in early embryos, where it is expressed in some cells of the frontal eye (Glardon et al., 1998; Kozmik et al., 2007; Vopalenski et al., 2012), in some cells of the ventral cerebral vesicle and in the lamellar body, a putative photoreceptive structure located in the dorsal roof of the cerebral vesicle (**Fig2G-J**). We found a very different pattern in the adult cerebral vesicle, with the ventral-anterior part now devoid of expression (**Fig2G-H and G’**) and the dorsal half showing expanded expression, thus resembling the situation in vertebrates. This dorsal *Pax4/6* domain is no longer associated with the lamellar body, which in adult brains is fragmented into scattered lamellar cells (Wicht and Lacalli 2005), but overlaps the *Emx* and *Lhx2/9* domains, as observed in the developing vertebrate telencephalon (Nat et al., 2013). In this overlapping region Pax4/6 reaches the anterior neural border, in a domain starting just above the neuropore (**Fig2I-J**). This position is clearly comparable to that of the Pax6 domain in the vertebrate telencephalon, also positioned at the most anterior-dorsal side of the brain. Contrasting *Pax4/6* expression in one-gill-slit embryos and in adults, we noted that in adults the ventral domain is posteriorly shifted towards the mid-caudal end of the cerebral vesicle. There, it extends caudally around the IO (**Fig2H**) and along the central canal passing through the primary motor centre (PMC), where a small number of *Pax4/6* expressing cells had already been observed in larvae (Vopalensky et al., 2012).

**Fig 2.**
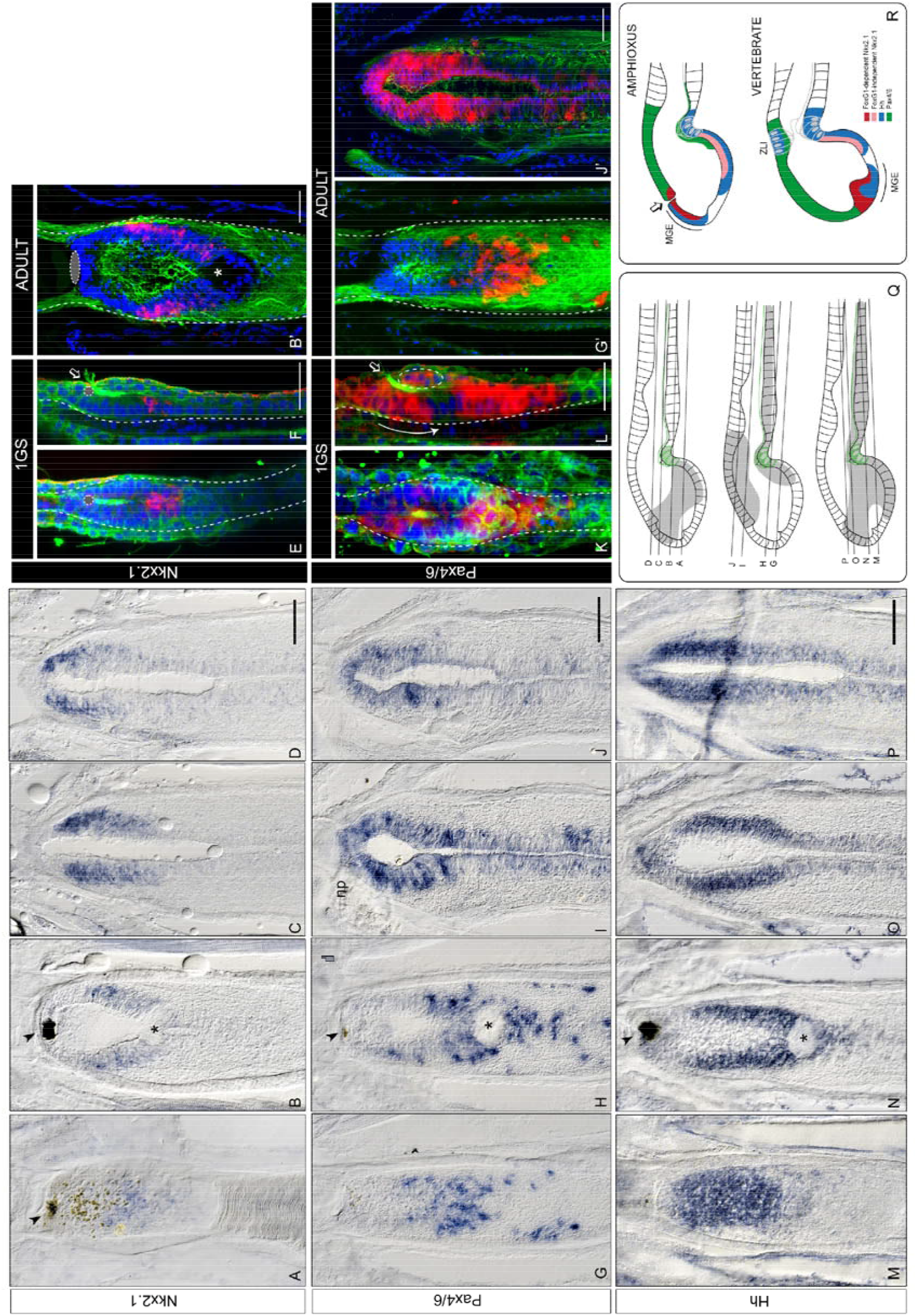
Late regionalisation of the telencephalic domain in amphioxus. Expression of *Nkx2.1* (**A-D**), *Pax4/6* (**G-J**) and *Hh* (**M-P**) is shown in coronal paraffin sections from ventral to dorsal with the ventricle of the cerebral vesicle (cv) centred in the images. *Nkx2.1* (**A-D**) is expanded anteriorly around the region between the pigment of the frontal eye and beneath the neuropore (**B-D**). This anterior expansion is especially visible when compared to the pattern in one-gill-slit larvae (**E-F**). **B’** is section **B** observed by confocal microscopy and counter-stained with acetylated tubulin (green) and dapi (blue). This is compared to a dorsal (**E**) and lateral (**F**) view of *Nkx2.1* expression in a one-gill-slit larva with the same counter-stain. *Nkx2.1* is noticeably ventral and posterior at the one-gill slit stage, closely localised around the anterior end of the IO (marked with asterisks in the adult(**B’**)). Note the strong acetylated tubulin staining for the luminal cilia exiting through the neuropore (empty arrow), clearly dividing the dorsal and ventral parts of the cv in one-gill-slit embryos. *Pax4/6* shifts posteriorly on the ventral side of the adult cv (**G-H**) and expands in the dorsal side (**I-J**). This is especially obvious when comparing the adult *Pax4/6* expression pattern to that of one-gill-slit embryos, shown in **K** and **L** as lateral and dorsal views, respectively, of a whole mount *in situ* imaged by confocal microscopy. The curved arrow in **L** indicates the direction shift of the *Pax4/6* ventral domain in comparison with **G’**(section in **G** imaged by confocal microscopy). The discontinuous circle in **L** indicates the very small dorsal *Pax4/6* domain in one-gill-slit embryos that is later expanded in adults, as seen in **J’**(section in **J** imaged by confocal microscopy).**M-P.** Hh expands anteriorly in the adult amphioxus brain. **Q**. Sectioning planes for *Nkx2.1* (**A-D**), *Pax4/6* (**G-J**) and *Hh* (**M-P**) on a lateral representation of an adult amphioxus brain. Grey shading indicates the expression domains for each gene (note that the anterior-most pigment of the frontal eye and the dorsal-most Joseph Cells Mantle are not represented in the diagrams for clarity). **R.** Overall the topography of these genes in adult amphioxus resembles that of early vertebrate embryos and further reveals the presence of an MGE region characterised by the overlapping expression of *FoxG1*-dependent-*Nkx2.1* (red) and Hh (blue). The *FoxG1*-independet-*Nk2.1* expression is represented in pink. Green represents the expression of *Pax6*, that in amphioxus has a ventral component unlike in vertebrates. The expression patterns of the vertebrate brain where drawn from expression patterns in the Allen Brain Atlas in mice embryos at E11.5. All scale bars in amphioxus sections are 50μm, except in **B’** and **G’-H’** that is 20μm as in **E-F** and **K-L**. Arrowheads indicate the pigment of the frontal eye in adults. The pigment of the frontal eye is otherwise indicated in the confocal images as discontinuous circles in grey. Discontinues lines show the outline of the brain in confocal images. Empty arrows indicate the anterior neuropore opening. All confocal images are counterstained with acetylated tubulin (shown in green) to label the axonal scaffold and with DAPI (shown in blue) to identify the nuclei of individual cells. Abbreviations: 1GS: One-gill-slit; np: neuropore.

In the developing vertebrate forebrain, *Nkx2.1* expression in the ventral telencephalon depends on *FoxG1*, whereas *Nkx2.1* hypothalamic expression does not (Martynoga et al., 2005; Danesin 2009). Since *Nkx2.1* is expressed in the absence of *FoxG1* in amphioxus embryos, it is thus likely that this domain is related to the vertebrate hypothalamus and not the vertebrate telencephalon. Consistent with these observations, we re-examined the expression of *Nkx2.1* in one-gill slit embryos, slightly later than the stages previously published in *B. lanceolatum* (Albuixech-Crespo et al., 2017), and observed that the *Nkx2.1* cells are indeed localised to the posterior-ventral side of the cerebral vesicle (**Fig2E-F**). Focusing on the adult amphioxus brain, we observed *Nkx2.1* expression in a new medial domain that, unlike the embryonic *Nkx2.1* domain extends to the anterior neural border of the vesicle (**Fig2A-D**) There, it partially overlaps with the *FoxG1* domain (**see Fig1D for comparison**), suggesting the existence of a *FoxG1*-dependent *Nkx2.1* domain, akin to the vertebrate telencephalon.

Expression of *Hh* has previously been reported in the neural plate and underlying notochord of neurulating amphioxus embryos. However, no expression has been detected in the cerebral vesicle or anterior neural plate at embryonic stages (Shimeld 1999, Shimeld et al 2007). In vertebrates, the expression of *Shh* in the anterior neural plate is required to induce the formation of the telencephalon (Retaux and Kano 2010). Consequently, the absence of *Hh* expression in the amphioxus cerebral vesicle during embryogenesis was previously interpreted as reflecting the absence of a telencephalon. In contrast, we found that in the adult amphioxus brain, *Hh* is strongly expressed in the cerebral vesicle, largely coinciding with *FoxG1* (**Fig2M-P**). In vertebrates, *Shh* and *FoxG1* also overlap and conditional removal of hedgehog signalling using a *FoxG1* driver results in a loss of ventral telencephalon patterning (Fuccillo et al., 2004). The overlapping expression of *FoxG1* and *Hh* in the anterior brain of the adult amphioxus suggests that a similar program could be in place in cephalochordates. Furthermore, our results show that the adult amphioxus brain has an intermediate *Hh+Nkx2.1* positive domain (**Fig2R**). This combinatorial code specifies the medial ganglionic eminence (MGE) in the vertebrate telencephalon (Sugahara et al., 2016). The MGE is the main source of dorsal telencephalic cells and cortical neurons and therefore it is essential for the formation of the dorsal telencephalon (Puelles et al., 1999; Marin and Rubenstein 2001; Moreno et al., 2009).

### Neuronal differentiation occurs late in the amphioxus brain

Since our results suggest that the adult amphioxus brain is laid out in a similar pattern to neural regions in early vertebrate embryos we next assessed the degree of differentiation of the amphioxus brain by examining the expression of markers of terminally differentiated neurons. Specifically, we examined the expression of transporters for acetylcholine and glutamate, V-AChT and V-GluT, used as markers for cholinergic and glutamatergic neurons respectively (**Fig3A-H**), as well as that of selected amphioxus neuropeptides (Jekely, 2013) (**Fig3I-Q**).

**Fig3.**
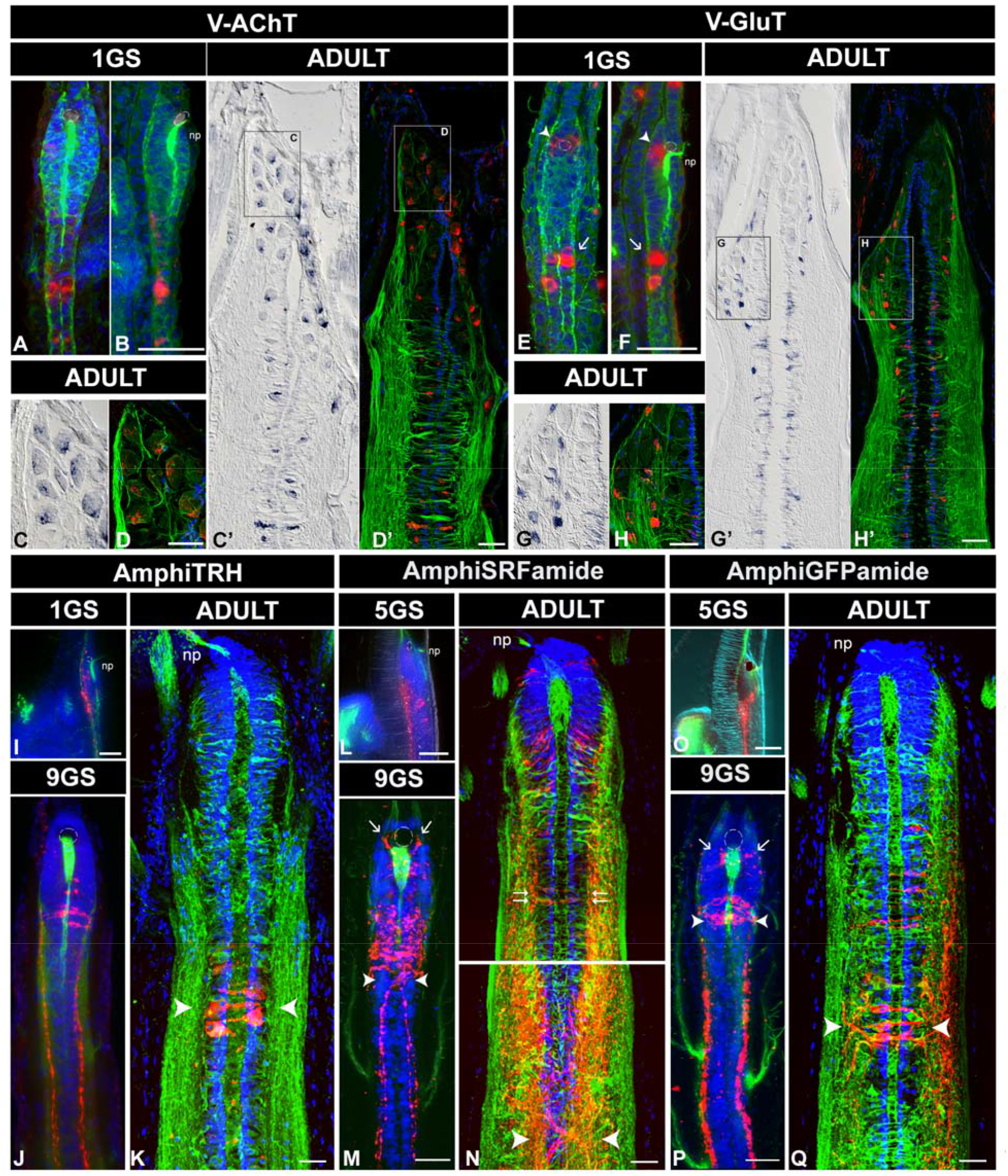
Late differentiating neuronal types in the amphioxus brain. A-D. Expression of the vesicular acetylcholine transporter (*V-AChT*) in whole mount *in situ* hybridisation of one-gill-slit embryos (**A-B**) and in coronal paraffin sections of adult brains (**C-D, C’-D’**). *V-AChT* is expressed in bilaterally arranged pairs of cells in the ventral side of the neural tube posterior to the cerebral vesicle. **A** and **B** are dorsal and lateral views, respectively, of the same embryo. **C** and **D** show magnified views of Joseph Cells expressing *V-AChT* in coronal paraffin sections of adult amphioxus brains (shown in full in **C’** and D’) under Nomarski (**C-C’**) and confocal microscopy (**D-D’**). **E-H.** Expression of the vesicular glutamate transporter (*V-GluT*) in whole mount *in situ* hybridisation of one-gill-slit embryos (**E-F**) and in coronal paraffin sections of adult brains (**G-H, G’-H’). E-F**. Dorsal and lateral views, respectively, of the same embryo, showing glutamatergic cells in association with the frontal eye (arrowheads) and posterior to the cv, again bilaterally paired and ventral. **G** and **H** show magnified views of putative lamellar cells expressing *V-GluT*, scattered amongst V-GluT negative Joseph Cells, in coronal paraffin sections of adult amphioxus brains (shown in full in **G’** and **H’**) under Nomarski (**G-G’**) and confocal microscopy (**H-H’**). **I-K.** Expression of AmphiTRH in 1GS (**I**-lateral view) and 9GS (**J**) embryos, and in adult vibratome sections (**K**). Arrowheads in **K** show the late developing transluminal cells. **L-N**. Expression of AmphiSRFamide in 5GS (**L**-lateral view) and 9GS (**M**) embryos, and in adult vibratome sections (**N**). In pre-metamorphic larvae (**M**) new AmphiSRFamide cells develop in association with the frontal eye (arrows) and the first contralateral descending projections are visible (arrowheads). These contralateral projections multiply in adults (arrowheads) and show the great amount of growth that the anterior brain of amphioxus undergoes as they locate farther away from the frontal eye/neuropore (np) region when compared to those of the pre-metamorphic larvae. Translumenal cells are indicated in **N** by double arrows. **O-Q**. Expression of AmphiGFPamide in 5GS (**O**-lateral view) and 9GS (**P**) embryos, and in adult vibratome sections (**Q**). Arrows in **P** show bilateral clusters of AmphiGFPamide cells caudal to the frontal eye and arrowheads point at the cluster of cells projecting to the dorsal midline. Arrowheads in **Q** show a later developing cluster of transluminal cells expressing AmphiGFPamide. Detailed views of **I, L** and **O** are shown in Supplementary Figure 2. All scale bars are 20μm. All images show dorsal views unless stated. All confocal images are counterstained with acetylated tubulin (shown in green) to label the axonal scaffold and with DAPI (shown in blue) to identify the nuclei of individual cells. Abbreviations: 1GS: one-gill-slit embryo; 5GS: five-gill-slits larva; 9GS: nine-gill-slits larva; np: neuropore.

First, and consistent with a late development of specific domains in the amphioxus cerebral vesicle, we observed that the dorsal roof of the adult brain is populated by cholinergic neurons (**see detail in Fig3C-D and Suppl. Fig2A-H**). These are not present in embryos, where the expression of V-AChT is instead ventral and mostly posterior to the cerebral vesicle (**Fig3A-B**). The dorsal cholinergic neurons are particular in size and morphology (see **Fig3C-D and SupplFig2A-H**), clearly identifiable as Joseph Cells (Wicht and Lacalli 2005).

Second, in mice 70-80% of cortical neurons (of telencephalic origin) are glutamatergic and their glutamatergic fate is set at their progenitor stage by the combinatorial action of *FoxG1, Lhx2, Pax6* and *Emx2* (Molyneaux et al., 2007). In amphioxus embryos few glutamatergic neurons are found only in the ventral-anterior floor of the cerebral vesicle, probably in association with the frontal eye (**Fig3E-F arrowheads**), and ventrally in the neural tube, with paired clusters starting at the level of the primary motor center (**Fig3E-F arrows**). In stark contrast, we have observed that adult amphioxus brains have several populations of glutamatergic neurons at different dorsoventral levels in the dorsal half of the cerebral vesicle (**Fig3G-H and G’-H’ and Suppl. Fig2I-P**). These comprise small spindle-shaped spinal fluid contacting (CSF-) neurons (**arrowheads in Suppl. Fig2M-N**), bilaterally arranged glutamatergic neurons deeply embedded in the neuropile (**arrows in Suppl. Fig2I-L**), and at the most dorsal level, scattered cells intermingling with the Joseph cells, probably corresponding to lamellate cells (**Suppl. Fig2O-P double arrows**) (see for comparison description by Castro et al., 2015). Our data suggest that similar to the vertebrate telencephalon, the cerebral vesicle in adult amphioxus is composed of several types of glutamatergic neurons. These are observed at the time when *FoxG1, Lhx2/9, Pax4/6* and *Emx* are expressed in the telencephalic region of amphioxus suggesting that some of these cells might be molecularly specified as in vertebrates.

By neuropeptide profiling we observed that also other neuronal types develop late. Using specific antibodies against three previously identified amphioxus neuropeptides, AmphiTRH, AmphiSRFamide and AmphiGFPamide (Jekely 2013) (**Fig3I-Q**), we saw that in general these begin to be expressed around the five-gill-slit stage (aprox. 3 months old animals) with the exception of AmphiTRH, which is already visible around the one-gill slit stage (**Fig3I-Q and Suppl. Fig3**). From pre-metamorphic stages until adulthood, the changing expression of these neuropeptides highlights the amount of late neuronal differentiation occurring in the brain of amphioxus, showing different neuronal types arising at particular times and locations within the brain at metamorphosis and beyond (Supplementary Note 1).

### The amphioxus brain grows slowly and outwards from the ventricular surface

Our data indicate that the amphioxus brain grows slowly and undergoes substantial remodelling during late development. To unravel the origin of cells newly added to the cerebral vesicle we focused on serotonergic cells that occur in the frontal eye and as two bilaterally arranged clusters posterior to the cerebral vesicle (Holland and Holland 1993, Lacalli et al., 1994, Moret et al., 2004). We were able to track these cells immunohistochemically through metamorphosis and we observed an increasing number of serotonergic cells in the frontal eye (**Fig4A-C**) and decussating serotonergic neurons (with axons crossing the midline) developing concomitantly with other decussating peptidergic neurons (**Fig4D-G**).

**Fig4.**
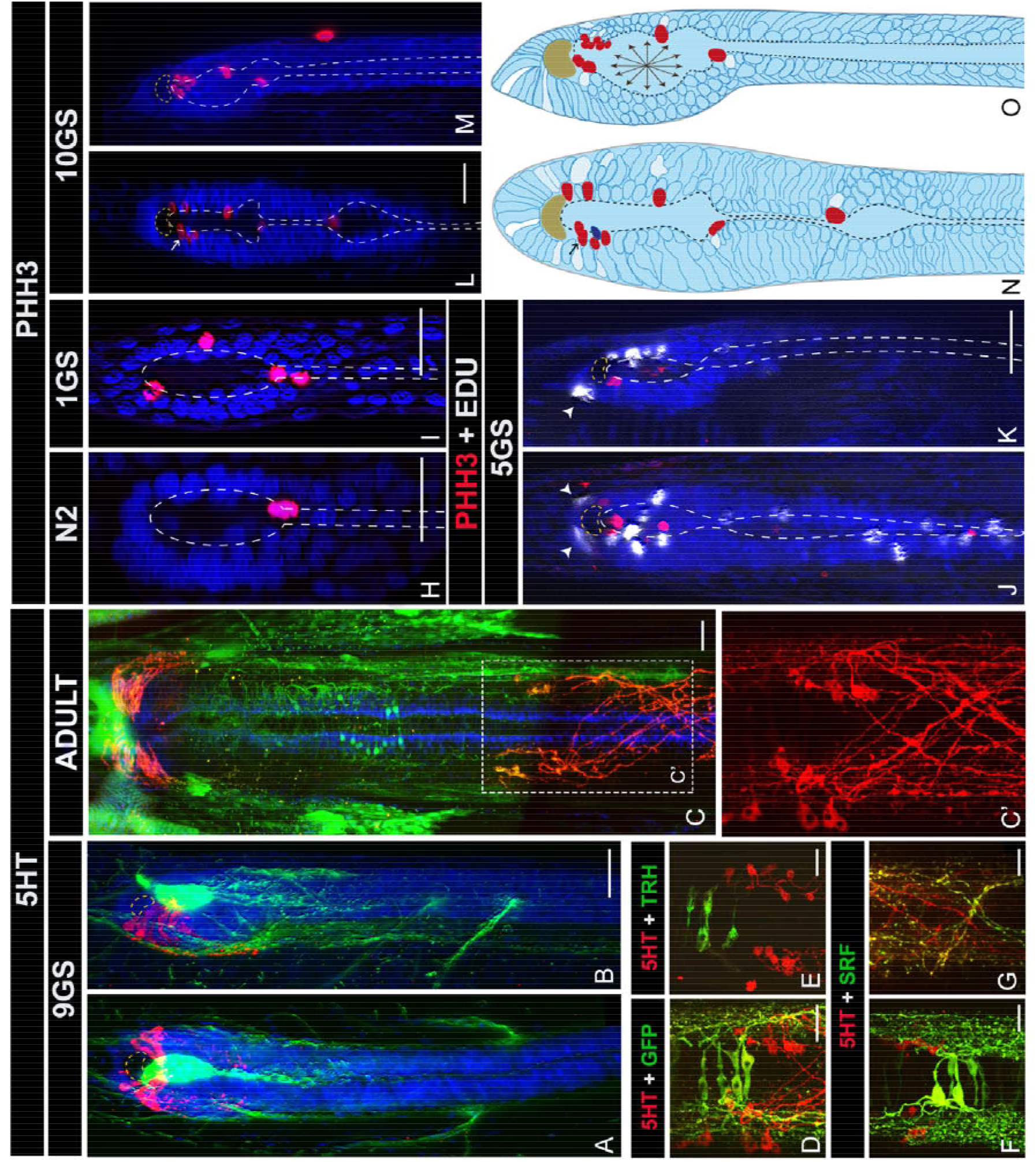
Late growth of the amphioxus brain. A-G. Expression of serotonin (5HT in red) is shown in pre-metamorphic larvae (9GS) in a dorsal (**A**) and lateral view (**B**). Five pairs of serotonin cells are organised around the pigment of the frontal eye (circled) at this stage. This anterior cluster of serotonin cells is bigger in the adult (**C**), showing around 16 pairs with greatly enlarged bodies. **C** shows the expression of serotonin in coronal vibratome sections of adult brains. In adults an additional cluster (previously denominated anterolateral cluster) appears caudally, framed in **C** and magnified in **C’**. The anterolateral cluster develops concomitantly with transluminal AmphiTRH, AmphiGFPamide and AmphiSRFamide cells, as shown by double immunohistochemistry in coronal vibratome sections of adult brains in **D-F**. **G** shows pairing of descending contralateral projections of serotonin and AmphiSRFamide cells in coronal vibratome sections of adult brains. **H-K**. Proliferation patterns in the amphioxus brain from neurula stages (**H**) to late pre-metamorphic embryos (**L-M**). At all stages mitotic cells (PHH3 in red) are localised in the ventricle (outlined) and only post-mitotic cells (white labelling in J and K) are pushed outwards away from the ventricle. Arrowheads in **J** and **K** indicate Edu + cells that divided in the ventricle and migrated basally. Occasionally it is possible to observe the apical-basal translocation of daughter cells, as indicated by arrows in **L** and **N**. **N-O** is our proposed model of brain growth in amphioxus, showing summarised distribution of PHH3 and Edu positive cells, as observed in different individuals in a dorsal (**N**) or lateral (**O**) view. Only a radial or centrifugal growth from the ventricle can explain the outwards distribution of Edu positive cells respect from the ventricular PHH3 positive cells in the amphioxus brain. The dark blue cell represents a progenitor cell that recently divided to give rise to a daughter cell still in red below it, also visible in **L**. All scale bars are 20μm. Abbreviations: N2: 7-8 somites neurula; 1GS: one-gill-slit embryo; 5GS: five-gill-slits larva; 9GS: nine-gill-slits larva; 10GS: ten-gill-slits larva.

To determine where the additional serotonergic cells were born in the cerebral vesicle, we next examined cell proliferation patterns in the brain of late developing embryos, as well as those of older metamorphic animals, via staining for the mitotic marker PHH3 (phoshorylated histone H3). We discovered mitotic cells covering the entire ventricular region at all stages, with PHH3+ cells being sparsely distributed over the ventricular surface (**Fig4H-M**). This suggests that neural stem cells are located and divide apically in the ventricular zone. Importantly, we did not observe PHH3+ cells at the anterior neural border, a site previously proposed as an anterior growth zone (Holland and Holland 2006), which suggests that cells move to the anterior neural border after dividing at the ventricular surface. The serotonergic cells added to the frontal eye are therefore added by intercalated growth from the ventricular surface.

As the cerebral vesicle grows, the number of visible mitotic bodies increases (**Fig4L-M**). Occasionally, we observed mitotic cells losing contact with the inner surface, suggesting that daughter cells translocate to the sub-ventricular zone, as observed in vertebrates (see arrow in **Fig4L** and **N**) (Guillemot 2005). To further track the fate of divided cells, we pulse-labelled late embryos with EdU (5-ethynyl-2’-deoxyuridine) two hours before fixation. As expected, this experiment located EdU positive cells basally to the ventricle, confirming that cells divide apically and translocate basally to sub-ventricular zones (**arrowheads in Fig4J-K**), as described in the vertebrate telencephalon. Throughout development, this centrifugal growth mode shapes and regionalises the cerebral vesicle (discontinuous line in **Fig4H-M, model in Fig4N-O ans Suppl. Fig4**). In this regard, we also uncovered here that the adult amphioxus brain is composed of three ventricles, with the main ventricle (**2v in SuppIFig4**) being further morphologically sub-divided into ventral and dorsal domains showing different cellular arrangements (**Suppl. Fig4D and E**).

## DISCUSSION

The assumed absence of a telencephalon homolog in the amphioxus cerebral vesicle has made the evolutionary origins of the vertebrate brain especially puzzling. Previous studies show the amphioxus brain as lacking any brain divisions, and more recently, it has been proposed to be a fused version of the vertebrate forebrain and midbrain devoid of any dorsal structures (which would include the telencephalic division of the forebrain) (Albuixech-Crespo et al., 2017). Our findings here provide compelling evidence for a telencephalon-like region in the anterior brain of amphioxus, which develops unexpectedly late. This telencephalic domain is subdivided into sub-regions of distinct molecular identity. Our results suggest a new evolutionary scenario where late activation of *Hh* in amphioxus induces the formation of a broad *FoxG1* domain (telencephalon) that further sub-divides into a dorsal domain expressing *Emx-Lhx2/9-Pax4/6*, a ventral domain expressing *Hh* and an intermediate *Hh-Nkx2.1*-positive domain, with the latter possibly representing a primitive MGE, which evolutionary origins were also controversial (Sugahara et al., 2017).

We have also uncovered the presence of dorsal populations of glutamatergic and cholinergic neurons, which in vertebrates are specified from neuronal progenitors located in the telencephalon. Our findings also indicate that these neuronal progenitors develop in amphioxus as in vertebrates, by proliferating in the ventricular zone and migrating basally to reach their final destination where they differentiate in situ. This is also here shown for the anterior serotonergic neurons, which development further suggests the amphioxus brain expands and regionalises by intercalated growth from the ventricular surface.

Herein, we provide substantial molecular and morphological evidence that there are two phases of brain development in amphioxus: i) an early, embryonic phase characterised by the development of ventral brain parts; ii) a late phase during which the dorsal parts of the brain develop, including a prominent counterpart of the vertebrate telencephalon. This late phase of development shapes the amphioxus brain into a layout unprecedently similar to that of early vertebrate brain.

Our findings make a strong case that the study of neurulating amphioxus embryos alone cannot reveal the extent of similarities between the amphioxus and the vertebrate brain. Therefore, previous notions that, for example, organiser regions would be missing in amphioxus (Pani et al, 2013; Albuixech-Crespo et al., 2017) will have to be re-evaluated. Also, our observations contrast with the traditional view that the amphioxus brain displays no signs of regionalisation other than the distribution of nerve exits. Overall, our data suggest that the adult amphioxus brain is a much better proxy for the vertebrate prosencephalon and thus encourage further cell type-specific comparisons across all life cycle-stages to finally solve the mysteries of chordate brain evolution.

## MATERIALS AND METHODS

### Collection, Breeding and Maintenace of Amphioxus

Wild catch collections of amphioxus were made in Banyuls-sur-mer (France) and in Kristineberg (Sweden). Once transported to the laboratory, the adult amphioxus were maintained, bred and the progeny raised as described in Benito-Gutierrez et al. 2013 in a custom made amphioxus facility.

### Gene Cloning

Sequences of interest were identified *in silico* using an unpublished transcriptome database generated in the laboratory. Primers were designed accordingly and used to amplify the transcripts of interest from a custom made SMARTer RACE (Clontech) cDNA library made from B.lanceolatum adult tissue. All genes were cloned in pCRII-TOPO TA vectors (Invitrogene, K4660-01) and *in vitro* transcribed using dual promoter sites for *in situ* probes. Probes were cleaned up using the RNeasy Kit (Quiagen, 74104).

### In situ hybridization and paraffin sectioning

For whole mount in situ hybridizations embryos were fixed and processed as in Holland 1999. For in situ hybridization on paraffin sections adult amphioxus were fixed in Neutral Buffered Formalin (HT501128, Sigma) for 24 hours. After fixation adults were dehydrated through an ascending series of methanol and further dissected in 100% methanol. The dissected heads were transferred to ethanol and then prepared for embedding in paraplast by xylene series: xylene:ethanol (1:1), xylene, xylene:paraplast (1:1) and finally paraplast (P3558, Sigma). Thereafter the prepared heads were transferred to a Leica Tissue Embedding Station (Leica EG1160). Paraffin blocks were sectioned using a Leica rotatory microtome (RM255) at a thickness of 12-14μm. The sections were left to float in a 37C DEPC treated water bath (Leica HI1210) and collected on SuperFrost Ultra Plus microscope slides (Menzel). The sections were flattened in a Leica flattening table (Leica HI1220) at 42C for approximately one hour. Sections were further fixed to the slides overnight at 37C. To prepare the slides for in situ hybridization, paraffin was dissolved by immersion in xylene. When sections looked clear they were rehydrated through a descending series of ethanol to PBS. The tissue was pretreated by immersion in HCl 0.2M for 15 minutes and later digested with 1μg/ml of Proteinase K at 37C for 40 minutes. The digestion was stopped by quick immersion in 0.2% glycine and glycine was thoroughly washed away with PBS. The subsequent hybridization steps were followed as for the in situs in wholemount embryos.

### Antibody Production and Immunohistochemestrv

For antibody production, rabbits were immunized with neuropeptides (sequences published by Jekely 2013) coupled to a carrier via an N-terminal Cys. Sera were affinity purified on a SulfoLink resin (Pierce) coupled to the Cys-containing peptides. The bound antibodies were washed extensively with PBS and with 0.5 M NaCl to remove weakly bound antibodies. Fractions were collected upon elution with 100 mM glycine, pH 2.7 and 2.3.

For immunohistochemistry in adult brains, adults were fixed as for paraffin sections in formalin for 24 hours. After that the brains were hydrated in PBS and embedded in 3% low melting agarose (A2790, Sigma). Agarose blocks were sectioned in a Leica Vibratome (VT 1200S) at a thickness of 60-80μm. Floating sections were collected in gelatin coated plates. The tissue was pre-treated as explained above for in situs and thoroughly washed afterwards in PBS-0.1%Triton-0.1%BSA. For both primary and secondary antibodies the tissue was blocked in PBS-0.1%Triton-5%NGS and incubated overnight at 4C with the appropriate antibody. All amphioxus neuropeptide antibodies were used at a concentration of 1μg/ml. Other antibodies were used as follows: anti-acetylated tubulin 1:250 (T 6793, Sigma); anti-5HT 1:200 (20080, Immunostar); anti-PHH3 1:200 (ab5176, Abcam). All secondary antibodies, goat anti-mouse-Alexa488 and goat anti-rabbit-Dylight549 (115-546-062 and 111-506-045, Jackson Immunoresearch) were used at 1:250.

For Edu detection embryos were incubated in seawater containing Edu at a concentration of 10μM for two hours before fixation. Embryos were harvested as for whole mount in situ hybridization and whole mount immunohistochemistry and fixed in 4%PFA-MOPS Buffer. Fluorescent detection of the incorporated Edu was performed following the manufacturer instructions (Click-iT^®^EdU Alexa Fluor^®^ 647 Imaging Kit, C10340, Invitrogen).

### Image Acquisition and Registration

Nomarski images were acquired with a Zeiss Imager M2 microscope. Confocal images were acquired with a Leica TCS SPE or a Leica TCS SP8. In all cases, sections and embryos were tile-scanned and thereafter processed using Deprecated Stiching Plugins (3D and others) in ImageJ (Fiji) (Preibisch et al., 2009).

